# H2O: A Foundation Model Bridging Histopathology to Spatial Multi-Omics Profiling

**DOI:** 10.64898/2026.04.21.717342

**Authors:** Yunjie Gu, Zihan Wu, Rui Yan, Zhikang Wang, Yuan Li, Senlin Lin, Yan Cui, Haoran Lai, Xin Luo, Shaohua Kevin Zhou, Zhiyuan Yuan, Jianhua Yao

**Author notes:** Corresponding Authors: Zihan Wu, Rui Yan, Shaohua Kevin Zhou, Zhiyuan Yuan and Jianhua Yao. These authors contributed equally (Yunjie Gu, Zihan Wu, Rui Yan).

## Abstract

Spatial omics technologies have revolutionized the molecular profiling of tissues but remain constrained by high costs and limited scalability. While hematoxylin and eosin (H&E) staining is ubiquitous, it lacks molecular specificity. Here, we present H2O (Histopathology to Omics), a generalist AI framework that bridges the modality gap between histopathology and spatial multi-omics, enabling the direct inference of spatial transcriptomics (ST) and proteomics (SP) landscapes from routine H&E images. H2O integrates Vision Transformers (ViT) with Large Language Models (LLM) via contrastive learning to align histological morphology with semantic molecular knowledge. This cross-modal approach allows the model to incorporate spatial expression profiles into histological pattern recognition, effectively decoding the molecular heterogeneity underlying tissue morphology. Trained on a pan-tissue dataset of 1.3 million paired H&E-spatial patches across 25 organs and cancer types, H2O predicts spatial omics expression from histology with high concordance to sequenced measurements and consistently outperforms state-of-the-art models across three cancer benchmarks. Notably, H2O recovers the *MIF-CD74/CD44* signaling axis directly from H&E images, highlighting its capacity to infer biologically meaningful cell-cell communication without molecular profiling. Applying on three additional public cohorts covering fetal and paediatric thymus tissues, human metastatic lymph node, and breast cancer, encompassing human development, 3D spatial frameworks, and integrative multi-omics, H2O yields biologically concordant insights, demonstrating superior accuracy, robustness, and generalizability across real-world applications in diverse scenarios. H2O converts routine histopathology into a portal for spatially resolved multi-omics profiling by computationally generating transcriptomic and proteomic landscapes, thereby enhancing tissue phenotyping and enabling scalable, integrative tissue-atlas construction.

## Introduction

Hematoxylin-and-eosin (H&E)-stained histology sections are routinely acquired and provides insights into disease mechanisms, playing a vital role in biomedical research and clinical practice [1,2,3,4]. However, H&E images are fundamentally limited by their inability to capture the molecular heterogeneity, specifically transcriptomic and proteomic landscapes, that drives disease progression. On the other hand, spatial omics profiling technologies provide spatially resolved mapping of transcripts, proteins and other molecules, which are crucial in revolutionizing our understanding from single biological events to the whole biological system. Emerging advancements in spatial omics technologies [5,6,7,8,9,10,11,12,13] have enabled high-resolution analysis of cell states, but their widespread clinical adoption is hindered by high cost, technical complexity, and incompatibility with archived formalin-fixed paraffin-embedded (FFPE) samples. There is an urgent need to bridge this gap by distilling high-dimensional molecular insights into routine histopathological workflows.

Recent advances in Vision Transformer (ViT) [14,15,16,17,18,19] architectures have catalyzed the development of a growing suite of general-purpose self-supervised models for computational histopathology, as represented by UNI [20], Virchow [21], CONCH [22], PLIP [23], mSTAR [24], and CTransPath [25]. In parallel, the adaptation of Large Language Model (LLM) [26,27,28,29,30,31] methodologies to the molecular life sciences has propelled the emergence of foundation models (FMs) capable of learning and interpreting the “languages” of biology, from genomic sequences to protein structures[32,33,34,35,36,37,38], such as scBERT [39], scGPT [35], scFoundation[40], scGPT-Spatial [41], and CellFM [42] for transcriptomics. Despite their individual successes, these models remain siloed. Vision FMs lack explicit biochemical context, while transcriptomics FMs lack spatial morphological grounding. Integrating these two modalities represents the next frontier in biological representation learning.

The rise of spatially resolved transcriptomics (ST) and proteomics (SP) has created a unique data landscape where high-resolution H&E images are inherently paired with molecular profiles. Early attempts to map the two modalities, like HisToGene [43], BLEEP [44], DeepPT [45] and OmiCLIP [46], relied on simple regression from image feature or retrieval mechanisms and failed to generalize across diverse biological contexts. Crucially, these methods ignore the pre-existing biological knowledge encoded in omics foundation models.

To address the existing challenge with the advanced FM technologies, we present H2O, a contrastive learning framework that aligns morphological FM with spatial omics guidance from ST FM to generate image embeddings with integrated histological and molecular information, enabling reliable prediction of spatially resolved molecular expression. To train H2O, we used 54,435,706 histopathology image patches from 44,992 samples and the HEST-1k [47] dataset, comprising 1.3 million data pairs from 1,229 paired ST and whole-slide image (ST-WSI) samples spanning 25 organs across multiple platforms. We verified the performance and generalizability of H2O on the previously withheld test samples of HEST-1k (ccRCC and PRAD) and an additional dataset (IDCLymphNode) for external validation.

To further assess the reliability and applicability of H2O, we conducted multi-scale validation across temporal, spatial, and modality dimensions using independent datasets including HTSA, OpenST and HTAPP (See Extended Data Fig. 1). H2O accurately recapitulated functional signaling networks, including the *MIF-CD44/CD74* axes essential for anti-apoptotic protection, and captured developmental trajectories, such as the postnatal activation of *CD19* and the fetal-to-pathological transition of *SPP1*. Beyond 2D patterns, 3D spatial omics constructions by H2O elucidated multicellular dynamics at tumor boundaries, revealing how *ACTA2*^+^ CAFs and *SPP1*^+^ macrophages trace invasive fronts in the lymph node parenchyma. Furthermore, the integration of H2O-predicted transcriptomics and proteomics refined tissue characterization by identifying unique tumor sub-regions that remained over-fragmented or obscured in single-modality profiling. These analyses consistently aligned H2O predictions with established biological principles, underscoring the framework’s capacity to derive novel mechanistic insights from computationally generated molecular maps.

## Results

### Overview of H2O as a cross-modal alignment method for histopathology and spatial omics

H2O is a multi-omics-guided histopathology foundation model designed to align whole-slide imaging features with spatially resolved transcriptomics and proteomics profiles through contrastive learning. To enable comprehensive coverage of various tissues, organs, and diseases (Extended Data Fig. 2) across both H&E and ST modalities, we curated a diverse dataset, which contains three major parts for H2O training and evaluation (Fig. 1a). Samples from The Cancer Genome Atlas (TCGA) [48], the Genotype-Tissue Expression (GTEx) [49] project, and additional in-house breast cancer collections (Methods) were used to train the histopathology FM. Then we employed a gene expression FM based on scGPT [35], initialized from whole-human pretrained checkpoints and further fine-tuned on ST data from the HEST-1k [47] dataset to encode spatial transcriptomics knowledge. Subsequently, we trained H2O using matched H&E and ST samples from the HEST-1k dataset. To infer transcriptomics profiles from a central image patch, H2O leverages morphological context from its local neighborhood and incorporates a Feature-wise Linear Modulation (FiLM) layer for resolution-aware feature calibration. See Extended Data Fig. 3 for ablation study of these modules. We further collected three additional datasets, HTSA, OpenST, and HTAPP, to investigate the ability of H2O in imparting molecular features to H&E features.

**Fig. 1.**
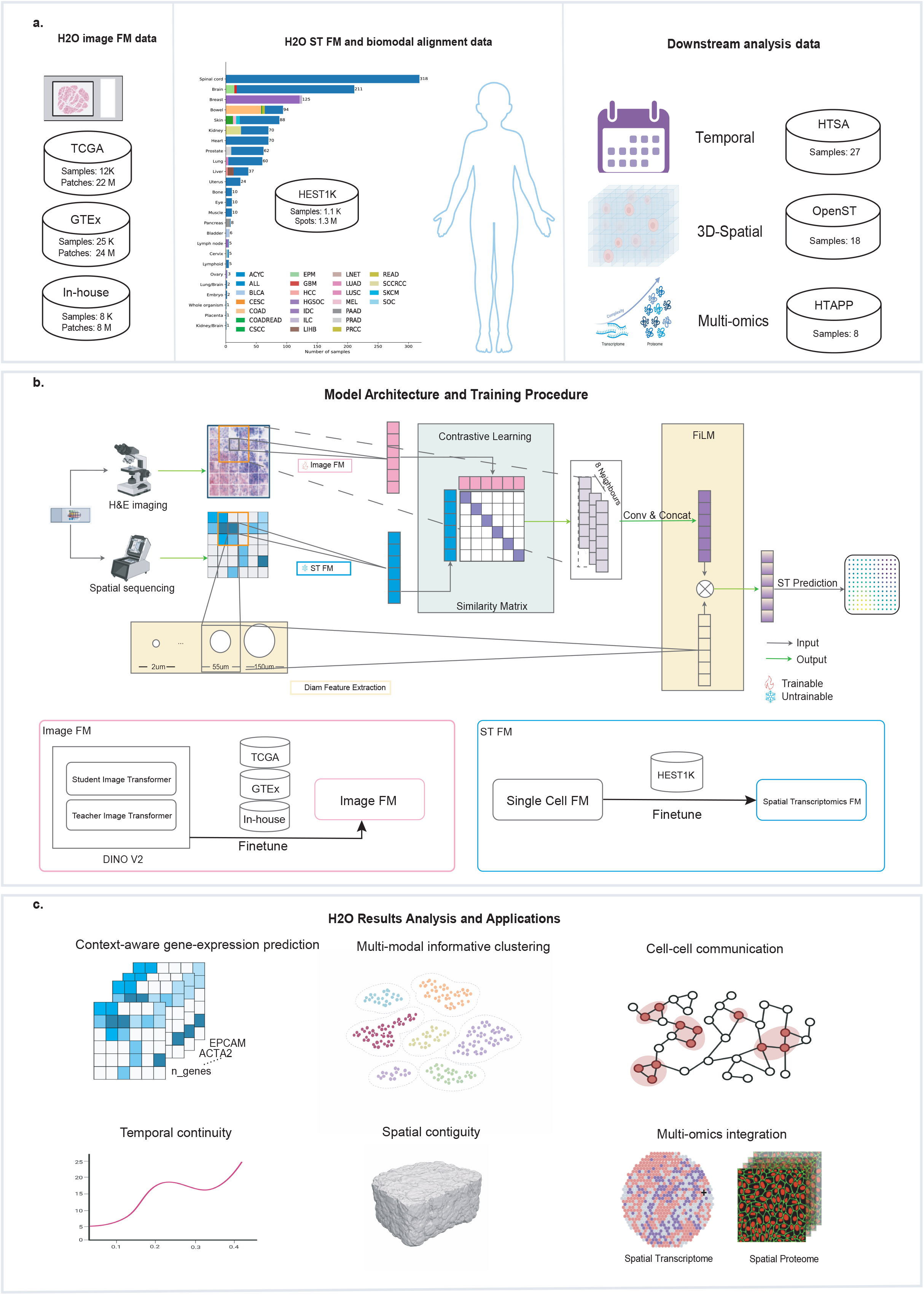
Training datasets, model architecture, and applications of H2O. **a**, Overview of datasets used in H2O. H&E images from TCGA, GTEx, and in-house breast cancer cohorts were used to train the histopathology image foundation model. Paired H&E-spatial transcriptomics data from HEST-1k were used for H2O model training. Independent datasets, including temporal thymus data from HTSA, 3D serial sections from OpenST, and multimodal transcriptomics-proteomics data from HTAPP, were used for downstream evaluation. **b**, Model architecture and training procedure. H2O comprises two steps: first, contrastive learning aligns a trainable histopathology foundation model and a parameter-frozen ST foundation model using paired image and gene expression data; second, grounded on the cross-modal aligned embeddings, the H2O decoder predicts the omics expression for each spot using the spatial context information. **c**, Downstream analyses enabled by H2O, including context-aware gene expression prediction, multi-modal informative clustering, cell-cell communication inference, temporal trajectory reconstruction, spatial contiguity analysis, multi-omics integration analysis.

#### A transcriptomics FM fine-tuned with ST data

Inspired by the great success of single-cell foundation models in representing the cellular characteristics, we utilized one of the representative methods, scGPT, to encode the transcriptomics features. In scGPT, the self-attention module learns intra-cellular gene-gene dependencies within a cell and inter-cellular interactions across different cells. Trained on 33 million single-cell transcriptomics data, representations from the model encode cell state and are sensitive to condition-specific variation, including developmental stages, disease progressions, and treatment responses.

However, a domain gap exists between ST and single-cell data. Further more, in comparison to single-cell data, ST exhibits greater heterogeneity owing to substantial platform variation, broadly spanning sequencing-based and imaging-based technologies [50]. Consequently, directly applying the scRNA-seq trained scGPT model to the ST data will inevitably induce improper representation of cells/spots. To address this, we fine-tuned the scGPT on the largest HEST-1k data, comprising 1.3 million cells/spots spanning sequencing-based platforms (10x Visium/Visium HD) and imaging-based assays (10x Xenium), enabling the learned representations to internalize cross-modal context while preserving modality-specific statistics. The fine-tuning process adapted the single-cell foundation model to spatial measurements and mitigated cross-modal domain shift relative to scRNA-seq while introducing ST knowledge.

#### An image FM aligned to ST latent space with contrastive learning

Recent work have proposed many histopathology foundation models specifically designed to extract morphology features from H&E images. Most of them are pre-trained on FFPE slices using self-supervised learning or multi-modal contrastive learning strategies. Nevertheless, the domain gap between FFPE and fresh frozen samples presents a significant challenge for histopathology foundation models. Most histopathology FMs are trained primarily on FFPE tissues, which are different from the fresh frozen samples typically used in ST technologies. To address this challenge, we first trained an in-house DINO-V2-based histopathology FM using FFPE and fresh frozen tissue samples to ensure its applicability to both tissue types. A total of 54,435,706 patches were extracted from 44,992 samples spanning 32 cancer types in The Cancer Genome Atlas (TCGA) and 40 tissue types from the Genotype-Tissue Expression (GTEx) dataset for model training. During patch extraction, rather than utilizing all available patches from each whole slide image (WSI), a checkerboard sampling strategy was employed, with a maximum of 2,000 patches selected per WSI to ensure representative diversity while mitigating spatial redundancy inherent in histopathology images. This selective sampling serves as a safeguard against training collapse and loss divergence, as excessive data redundancy in large-model pre-training has been shown to compromise both convergence stability and downstream generalizability [51].

Following this, we employed a contrastive learning framework to align H&E image embeddings with the latent space of ST by maximizing similarity between positive H&E-ST pairs while simultaneously minimizing similarity between negative pairs profiles. This alignment encourages the image encoder to learn characteristics from both the histopathology images and transcriptomics profiles, preserving morphology characteristics while introducing potential omics knowledge. Crucially, it distills intra- and inter-cellular information from transcriptomics FM, such as gene-gene dependencies and neighborhood interactions, into the image representation, improving biological fidelity and robustness across platforms.

Predicting ST profiles using fixed-size image patches fails to accommodate the intrinsic variability across different platforms and resolutions. To retain this information, we adopted diameter-conditioned learning, where the encoder was conditioned on spot size, providing platform- and resolution-aware context during training to improve the coupling between image features and expression magnitude and enhance cross-platform transferability. And to improve quantitative awareness in prediction, we adopted a FiLM module to integrate auxiliary calibration information.

#### The H2O framework

In summary, H2O establishes a foundational cross-modal framework that anchors histopathological features into a high-dimensional molecular space, enabling the direct translation of standard H&E images into rich, molecularly informed landscapes (Fig. 1b). By leveraging contrastive alignment and platform-specific conditioning, H2O transcends simple morphological correlations to capture the latent biological programs linking tissue architecture to gene and protein expression. This versatile representation facilitates a broad spectrum of downstream tasks (Fig. 1c), ranging from context-aware expression prediction and multi-modal clustering to the elucidation of cell-cell communication, temporal trajectories, and 3D spatial contiguity. Ultimately, H2O provides a scalable pipeline to unlock deep biological insights and multi-omics integration from existing histological archives.

### Evaluation of H2O for spatial transcriptomics prediction from H&E images

Accurately predicting ST from H&E images holds great significance in efficient and economical biology analysis. To assess the predictive capacity of H2O, we evaluated it on 53 test samples from the HEST-1k dataset that had been withheld from both the scGPT fine-tuning and contrastive learning phases. We benchmarked H2O against HisToGene [43], BLEEP [44], and DeepPT [45], and OmiCLIP [46], by training and testing all methods on the clear cell renal cell carcinoma (kidney cancer, ccRCC, 24 samples) and prostate adenocarcinoma (prostate cancer, PRAD, 23 samples) data of HEST-1k, predicting 5,033 global highly variable genes (HVGs) of this dataset. Across both benchmarks, H2O consistently outperformed the competitive methods in terms of Pearson Correlation Coefficient (PCC) and Spearman Rank Correlation Coefficient (SRCC) (Fig. 2a). It is evident that H2O holds an obvious advantage over other state-of-the-art methods across the evaluation metrics.

**Fig. 2.**
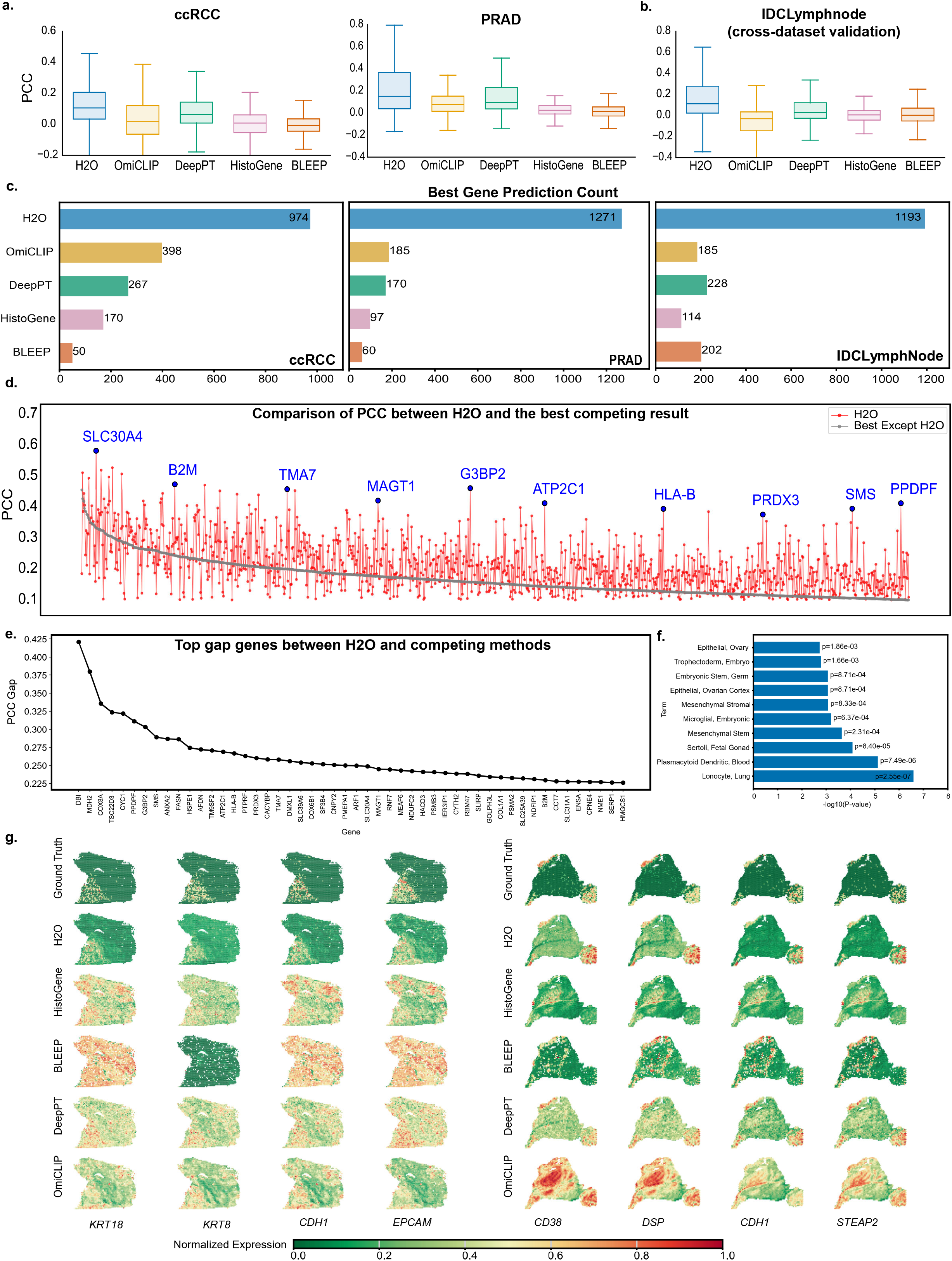
Evaluating H2O for spatial transcriptomics prediction from H&E images. **a**, Performance of H2O and compared methods on ccRCC and PRAD datasets in terms of PCC and SPCC. **b**, Generalization benchmarking on the independent IDCLymphNode dataset. **c**, Number of genes for which each method achieved the top correlation with ground truth in ccRCC, PRAD, and IDCLymphNode test samples. **d**, Comparison of PCC for all genes between H2O and the best competing result. Genes with a PCC less than 0.1 in either method were not shown. H2O yielded higher correlations in predicting most genes. Blue marks point out genes with significant PCC gaps. See Extended Table for details of gene symbols. **e**, Top 100 genes with the largest gaps in PCC of H2O and the mean value of competing methods. These genes gaps illustrate where transcriptomic priors provided decisive advantages beyond morphology alone. **f**, Functional enrichment of the most predictable genes by H2O using gene set from GSEA. **g**, Spatial maps of selected marker genes comparing ground truth (GT) with predictions from H2O, HistoGene, BLEEP, DeepPT and OmiCLIP.

To further test the models, we applied the well-trained prediction methods on HEST-1k-ccRCC and HEST-1k-PRAD to an independent test dataset of axillary lymph nodes of invasive ductal carcinoma, IDCLymphNode (Fig. 2b). In this comparison, H2O still holds advantages over other methods, demonstrating its strong capability in cross-biological scenes, i.e., different cancer types. We attribute this to the utilization of scGPT in the training phrase, which highlights that molecular knowledge encoded by scGPT-FT is transferable across cancer types, providing H2O with a strong inductive bias for cross-dataset generalization. The leading performance of H2O across all three benchmarks underscored its robustness across the transcriptome.

Predicting gene expressions from histology is fundamentally challenging. For such predictions to support meaningful downstream biological interpretation, the model must reliably recover a core set of functionally relevant gene, including established cell-type markers and pathway-essential genes. We evaluate the efficacy of H2O alongside traditional benchmarks, specifically examining its performance across selected gene sets to better handle increased data complexity. We first selected a fixed global set of 2,000 genes by ranking mean PCC across all samples for each dataset. Then, for each model, we calculated the number of genes for which the model achieved the highest average PCC across all samples (Fig. 2c). H2O achieved the largest share of the best-performing genes, underscoring its robustness across the transcriptome. We further contrasted the performance of each single gene predicted by H2O and the best prediction of other competing methods (including DeepPT, HistoGene, OmiCLIP and BLEEP). As shown in Fig. 2d, H2O achieved higher correlations across most of these genes than other methods, reinforcing that supervision from a transcriptomics foundation model yields substantial gains in per-gene predictive performance. We next identified genes showing the largest performance gaps between H2O and other methods, which indicated those benefiting most from incorporating transcriptomic priors (Fig. 2e). Functional enrichment of these genes indicated strong associations with established biological pathways [52,53], reinforcing the utility of H2O for high-fidelity molecular and microenvironmental characterization (Fig. 2e). Spatial maps of marker genes *(KRT18, KRT8, CDH1, EPCAM in sample 1 and CD38, DSP, CDH1, STEAP2 in sample 2*) samples of the benchmark set PRAD, showed that H2O not only more faithfully generated spatial gene distributions than the best non-prior methods, DeepPT, but also defeat the omics-prior methods, BLEEP and OmiCLIP (Fig. 2f). We also explored H2O’s prediction visualization of top 6 Best Predicted Genes, Highly Variable Genes and Spatial Variable Genes from the test samples (Extended Data Fig. 4-6).

Through comprehensive benchmarking, cross-dataset validation, and gene-level quantitative and visual analyses, our results showed that H2O demonstrated superior fidelity in recovering fine-scale and biologically meaningful spatial expression patterns compared with existing methods.

### Validation of H2O ability to reveals molecular expression features and functional signaling patterns

The reliability of downstream analysis is crucial for ST data. In this section, we aimed to utilize the H2O-predicted ST data for several downstream analyses, verifying the utility of the well-trained H2O prediction and identifying potential biological insights infeasible from the H&E histopathology images.

One central downstream task is spatial domain recognition, which requires the image-omics model predictions can reliably reproduce the molecular structure of the underlying tissue. To verify that H2O recapitulates experimentally validated expression patterns, we first applied k-means clustering across varying values of k. Clusters derived from k-means clustering after principal component analysis (PCA) on H2O-predicted expression more closely resembled clusters identified from experimental profiles than those obtained directly from image embeddings (Fig. 3a). This proves that H2O recovers molecular information absent from raw histology, leading to predicted expression profiles that more closely match experiment-measured patterns. Quantitatively, Adjusted Rand Index (ARI) and Adjusted Mutual Information (AMI) scores confirmed the superiority of H2O compared against image-only embeddings, with values of 0.66 vs. 0.34 (ARI) and 0.63 vs. 0.44 (AMI), respectively (Fig. 3b). These results demonstrated that transcriptome-informed predictions capture biologically relevant structures beyond what can be inferred from morphology alone.

**Fig. 3.**
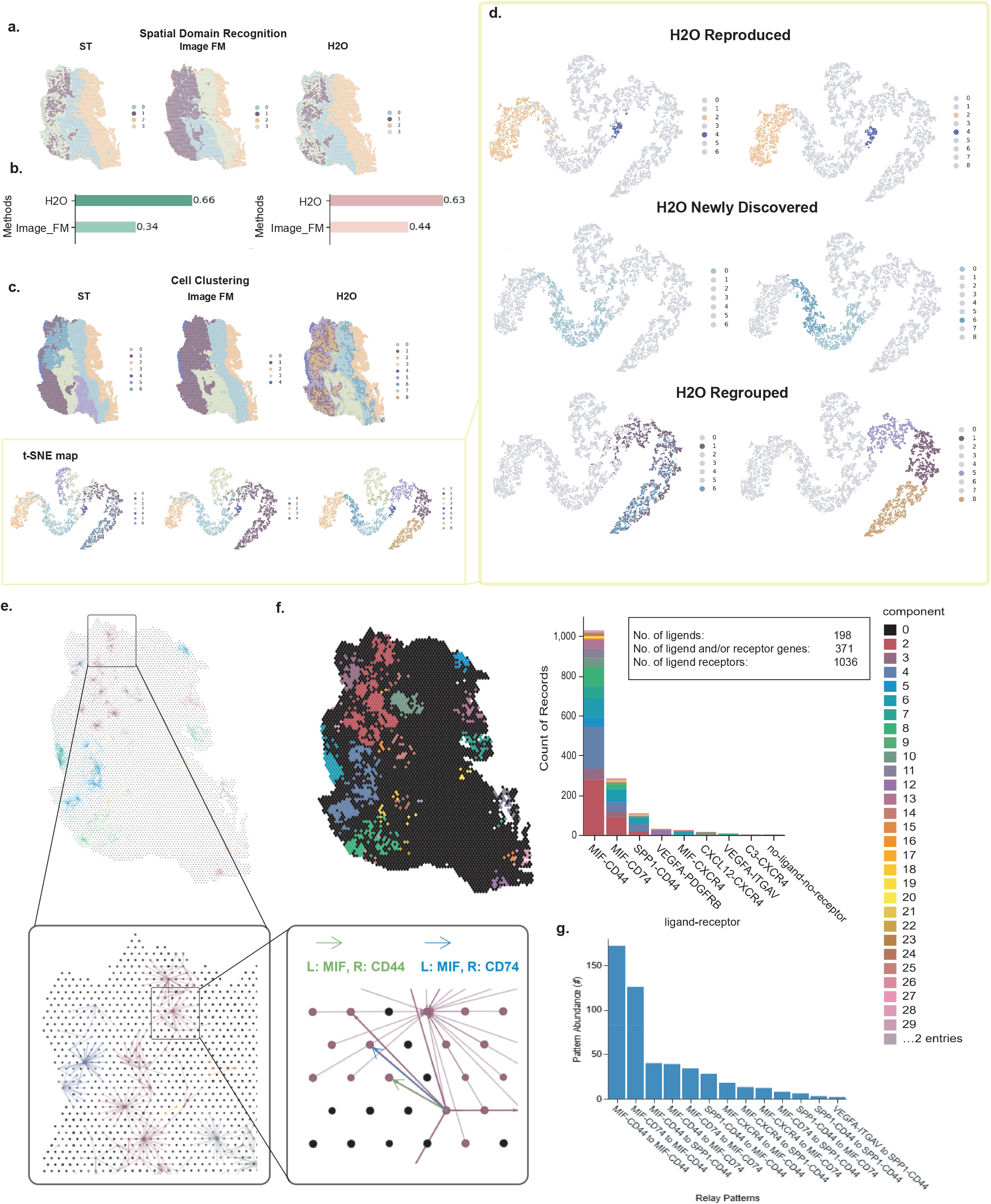
Validation of H2O predictions through clustering and cell-cell communication analysis. **a**, Comparison of spatial domain recognition derived by k-means clustering on experiment-derived ST profiles (ST), morphology-based embeddings produced by the unaligned image FM of H2O (Image FM), and H2O-predicted profiles (H2O). **b**, Quantitative evaluation of k-means clustering similarity at k=4. Adjusted Rand Index (ARI) and Adjusted Mutual Information (AMI) scores showed H2O (0.66 ARI, 0.63 AMI) substantially outperformed Image FM embeddings (0.34 ARI, 0.44 AMI). **c**, Upper: Leiden clustering applied to ST, Image FM embeddings, and H2O prediction. Lower: t-SNE plot of ST, Image FM embeddings and H2O Prediction colored by Leiden clusters; **d**, Categorization of Leiden clusters. H2O-derived clusters were classified as reproduced, newly discovered, or regrouped relative to the ST results, demonstrating the additional value of transcriptome-informed embeddings. **e**, Cell-cell interaction analysis using CellNEST on H2O-predicted ST profiles. Predicted spatial expression enabled inference of local signaling neighborhoods, with detailed network visualization of ligand-receptor connectivity. Highlighted interactions included MIF-CD44 and MIF-CD74, both are crucial for MIF’s role in protecting against apoptosis. **f**, Left: Spatial localization of predicted communication networks mapped onto tissue context inferenced from H2O predictions. Right: Frequency of inferred ligand-receptor pairs across clusters, with MIF-CD44 and MIF-CD74 emerging as dominant interactions. **g**, Relay pattern analysis of ligand-receptor cascades, showing dominant interaction motifs, including MIF-centered pathways.

We next sought to assess how the joint embeddings of H2O improve the cell clustering task. Inspired by the MUSE [54] framework, we evaluated multimodal latent spaces by categorizing clusters as reproduced, newly discovered, or regrouped relative to unimodal results. Specifically, we compared Leiden clusters derived from image features alone with those obtained from the joint latent space provided by H2O (Fig. 3c). This analysis revealed that transcriptome-guided embeddings not only captured known histological structures but also enabled the uncovering of new or reorganized clusters that were not apparent in image-only space (Fig. 3d). Together, these analyses, spanning both target-based k-means and resolution-based Leiden clustering, demonstrated that H2O effectively transferred molecular information into image embeddings through the transcriptome-guided contrastive training procedure of the image encoder.

To further validate the reliability of H2O-predicted gene expressions, we turned to cell-cell communication, which reflects higher-order tissue architecture and plays a pivotal role in processes such as immune modulation, tumor progression, and therapeutic response. Taking the predicted gene-expression data as input, the cell-cell commutation method CellNEST [55] identified in total 1,036 ligand-receptor interactions (Fig. 3e-f) and 30 densely communicating regions (Fig. 3f). Among these interactions, *MIF-CD44* and *MIF-CD74* emerged as the two most prominent pairs (Fig. 3f). Further exploration of the cascade abundance of the two ligand-receptor pairs showed the *MIF-CD74* to *MIF-CD74* pair and *MIF-CD74* to *MIF-CD44* pair are highly represented (Fig. 3g). These findings are consistent with prior studies [56] showing that *CD74* and *CD44*, either independently or as heterodimeric complexes, are essential for MIF-mediated protection against apoptosis. Recapitulating such well-established molecular interactions indicates that H2O not only reproduces single-gene expression patterns but also preserves functional signaling relationships, highlighting its value for precise molecular and microenvironmental profiling. Heatmap matrices among selective genes of ST and H2O predictions shows similar patterns proves this knowledge-aware learning, see details in Extended Data Fig. 7.

### H2O generalizes to developmental systems in the human thymus

A fundamental question for an image-based molecular predictor is whether its outputs reflect genuine molecular signatures and remain consistent with prior biological knowledge. To test the reliability, we applied H2O to the Human Thymus Spatial Atlas (HTSA) [57] (Fig. 4a). HTSA provides a longitudinal series capturing T cell development from fetal through early postnatal life, containing 28 samples across a timeline spanning from 13 weeks post-fertilization to 2 years of age, with high-resolution multiplexed imaging, spatial transcriptomics, and matched single-cell transcriptomics at each time point. This systematic temporal coverage makes the dataset ideally suited to assess whether morphology-based predictions can reconstruct both immune developmental trajectories and spatial microenvironment dynamics across maturation. The availability of histopathology images aligned to spatial omics data provided an ideal testing ground to assess whether H2O could reconstruct developmental processes directly from routine H&E slides.

**Fig. 4.**
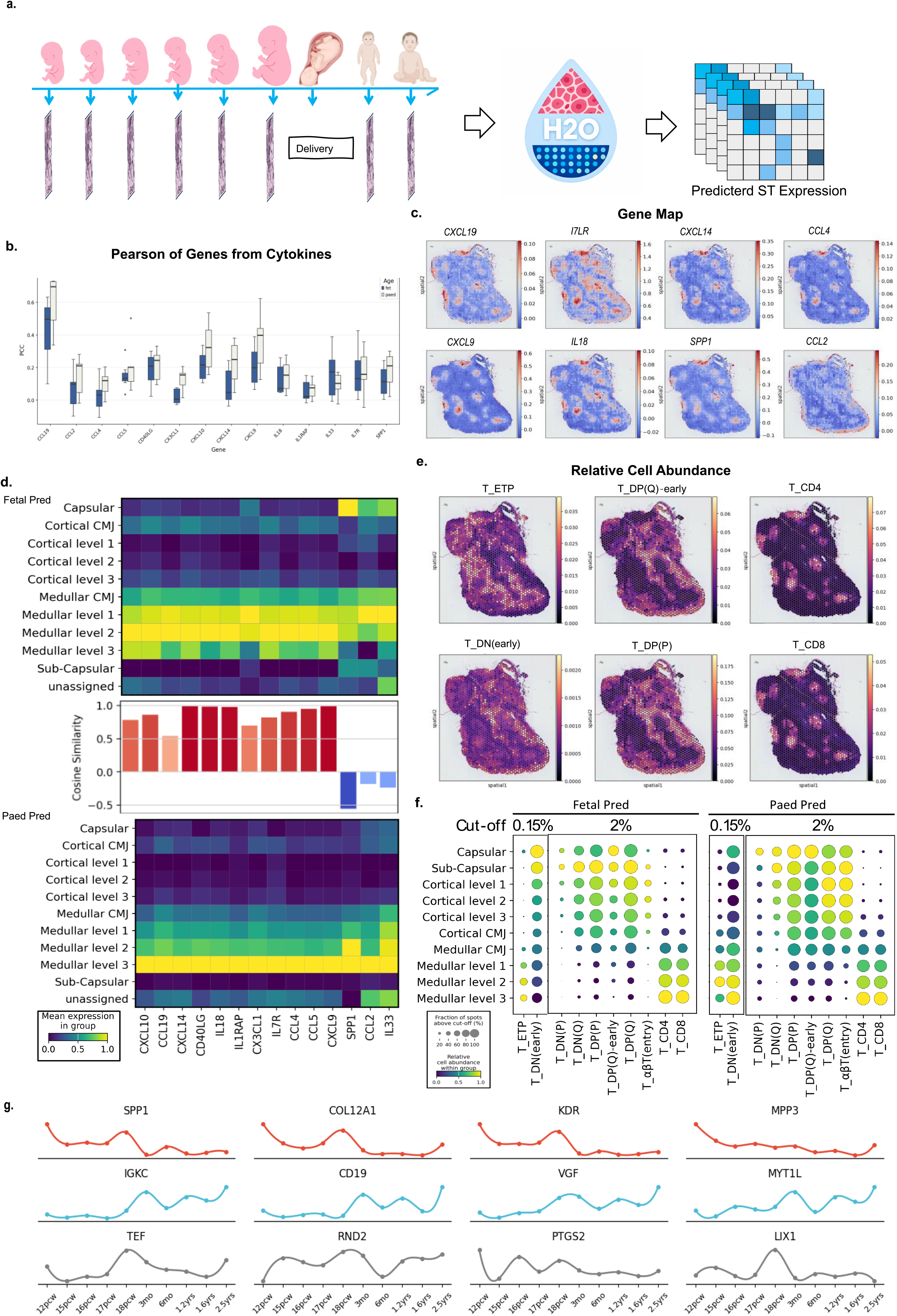
H2O generalizes to developmental systems in the human thymus. **a**, Overview of the H2O application on the Human Thymus Cell Atlas. Histopathology images of developmental stages from fetal to paediatric thymus were processed by H2O to predict ST across time. **b**, Prediction evaluation for cytokine genes. Boxplots show Pearson Correlation Coefficients (PCCs) between H2O predictions and experiment-derived profiles, stratified by developmental stage. **c**, Spatial expression maps of representative cytokines marker genes. **d**, Heatmap of spatial region distribution of representative T-cell marker genes at fetus and paediatric stages. The top panel belongs to the fetus and the bottom one belongs to the paediatric state. Middle is cosine similarity of the gene distributions between fetal and paediatric stages. Most markers displayed conserved spatial patterns, whereas SPP1, CCL2, and IL33 showed stage-dependent redistribution from the capsular region in fetal thymus to the medulla in paediatric thymus. **e**, Relative cell abundance (cell proportion) of cell-type deconvolution results from H2O prediction using Cell2location; Sub-types of T cells representing crucial stages in T-cell development are shown. **f**, Regional distribution of the cell-type deconvolution results from H2O-predicted ST data. Dot plots depict T-cell subtype enrichment across thymic compartments, highlighting stage-specific shifts of the rare ETP and DN(early) stages. Size of the dots represents the fraction of spots larger than the 0.15% or 2% cut-off; Color of the dots represents the mean relative cell abundance. **g**, Gene expression trends of H2O-predicted data across developmental stages, each panel shows the temporal dynamics of a specific cytokine- or immune-related gene during thymic development from early gestation to postnatal stages.

The HTSA dataset provides a continuous, scale- and rotation-invariant morphological common coordinate framework (CCF) for the thymus, enabling systematic comparisons across donors and stages. Leveraging this framework, we stratified H2O predictions into fetal and paediatric groups and quantified the expression of canonical T-cell marker genes of the two groups (Fig. 4b). Fig.4c (more in Extended Data Fig. 8) shows the spatial distribution of 8 H2O-predicted marker genes (*CXCL19, IL7R, CXCL14, CCL4, CXCL9, IL18, SPP1, CCL2*). Correlating these distributions across developmental stages revealed that most cytokine genes maintained highly conserved spatial organization, consistent with the notion that the thymic microenvironment remains functionally stable during T-cell maturation [57] (Fig. 4d). Importantly, three genes, *SPP1, CCL2*, and *IL33*, showed substantially lower cosine similarities between fetal and paediatric stages, suggesting developmental stage-dependent reorganization. Mapping these genes back to the CCF indicated a redistribution of expressions from the capsular region in the fetal thymus to the medullary compartment in the paediatric thymus, reflecting a shift in microenvironmental niches as development progresses (Fig. 4d).

To disentangle the cellular basis of these shifts, we performed cell type deconvolution with Cell2location [58] on the H2O-predicted ST profiles using matched scRNA-seq data of the same tissue type as reference (Fig. 4e,f). The deconvolution results enabled us to estimate the average spatial distribution of cell populations along the CMA axes. To filter out cell types with low abundance, we applied cut-offs of 0.15% and 2%, representing the minimum relative cell abundance required for spot inclusion. The results are visualized in Fig. 4f, where the spot size corresponds to the fraction of spots exceeding the abundance cut-off and the spot color indicates the mean relative abundance of each cell type in each region. This analysis revealed that stage-specific expression differences were largely attributable to changes in Early T-cell Progenitors (T-ETP) and the broader Early Double-Negative stage of T cells (T_DN(early)) in the medulla in both fetal and paediatric tissue, aligning with previous reports highlighting the role of stromal and immune niche remodeling in shaping T-cell differentiation [59]. For example, fibroblasts and endothelial cells are known to contribute to cytokine gradients, while TECs form the structural basis of positive and negative selection. Macrophages, in turn, regulate apoptotic clearance during thymocyte turnover, providing a functional explanation for their redistribution. Thus, by aligning cell type abundance to the predicted ST spots, H2O reconstructed the spatial trajectory of T-lineage differentiation, revealing both conserved programs and stage-dependent remodeling of the thymic microenvironment (Fig. 4f). Importantly, the recovered cytokine maps recapitulated established paradigms of thymic development, where T-cell migration cues remain broadly conserved across fetal and paediatric stages while stromal and innate immune components undergo localized reorganization.

To further validate the biological fidelity of H2O, we analyzed the developmental trajectories of key genes predicted by H2O, observing expression patterns highly consistent with established knowledge (Fig. 4g). Specifically, the B-cell coreceptor *CD19* was nearly undetectable during fetal stages but significantly upregulated after birth. This reflects a “silent-in-fetus, increased-after-birth” pattern, which is fundamentally aligned with the ontogeny of the human adaptive immune system [60]. In contrast, SPP1 (Osteopontin) was highly expressed during fetal development, where it orchestrated cell adhesion, migration, and tissue mineralization essential for organogenesis [61], declined to low levels postnatally, and was pathologically re-activated in cancer and inflammatory contexts [62]. Given its widespread re-expression across malignancies, SPP1 serves as a general cancer marker, a premise we further investigate in the following section.

These results show that H2O not only possesses the capability of predicting ST across cancer contexts but also demonstrates practical application in developmental systems with robust generalizability, demonstrating the potential of H2O to be used as a low-cost, image-based approach to uncover both stable developmental blueprints and dynamic cellular transitions in complex tissues.

### H2O constructs 3D architecture to explore tumor dynamics

Deciphering the 3D architecture of tumor tissue, e.g., tumor invasion, immune infiltration, and stromal modeling, is critical to understanding the tumor biology and characterize the patients. Nevertheless, existing 3D ST methods are significantly limited by complex experiment conditions, low sensitivity, ,small panel size and high cost[63]. Translating spatial omics from routinely accessible H&E images provides great chance.

To address this challenge, we utilized the OpenST dataset [63], comprising 30 serial slices (19 ST-H&E pairs and 11 H&E images) from a human metastatic lymph node, to evaluate H2O’s capacity for cost-effective 3D construction. We first registered the serial sessions to construct a virtual three-dimensional tissue block. Then we implemented H2O to these H&E images, predicting in situ transcriptomics data (Fig. 5a).

**Fig. 5.**
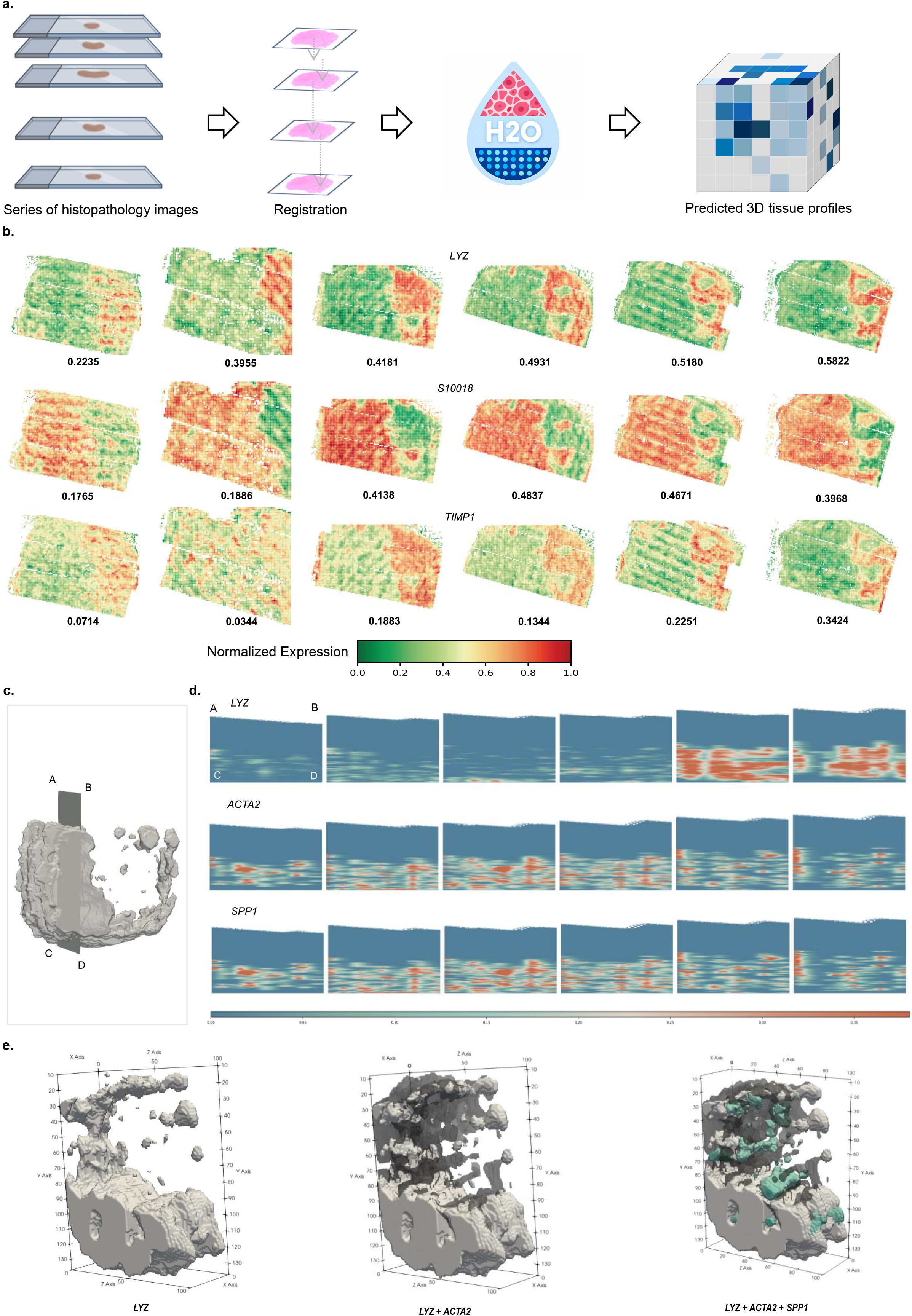
H2O constructs 3D tumor architecture with clinical translational potential. **a**, Overview of the workflow for applying H2O to the OpenST dataset. Serial H&E slides from a metastatic lymph node were aligned and processed with H2O to predict spatial gene expression across sections, allowing a three-dimensional construction. **b**, H2O-predicted expression of selected marker genes across sections, including TIMP1 for cancer-associated fibroblasts, LYZ for macrophages, and S100A8 for lymph node-associated myeloid cells. PCC between prediction and experiments measured profiles are shown at the left bottom corner of each sample. Predictions captured both the compartmentalization of lymph node tissue and tumor-associated infiltration. **c**, schematic of 3D sections. **d**, H2O-predicted cross-sectional views of marker genes, including LYZ, ACTA2 and SPP1, in the 3D stack. **e**, Three-dimensional renderings of the marker expression of H2O predictions. The visualization revealed continuous tumor-stroma structures and invasive patterns of tumor penetration into the lymph node, which cannot be resolved from individual two-dimensional sections.

We examined section-level predictions of canonical marker genes representing key cellular compartments, including *LYZ* for lymph node-associated macrophages, *S100A8* for inflammatory myeloid cells, and *TIMP1* for fibroblasts (CAMs) [63]. H2O-predicted expression patterns showed high concordance with experiment-derived distributions, providing confidence for subsequent 3D integration (Fig. 5b). We then stacked the predictions in sequential section order and visualized the resulting volume to examine the spatial distribution of marker genes along the z-axis. We presented z-axis cross-sections at representative x-axis positions to highlight local variations in the organization of marker genes across the tissue (Fig. 5c,d). Our results demonstrated that H2O captured marker enrichment in tumor cores representing by SPP1, stromal boundaries enriched with ACTA2, and lymph node sinuses abundant with LYZ, consistent with established spatial biology conclusions of metastatic progression [63].

Moreover, using the 3D rendering tool [64], we further visualized the predicted omics features in the real 3D space (Fig. 5e). These 3D reconstructions elucidated how CAFs (*ACTA2*^+^) and cancer-associated macrophages (*SPP1*^+^) distinctively delineate tumor boundaries and trace invasive fronts as malignant cells penetrate the *LYZ*-rich lymph node parenchyma. Importantly, the 3D perspective uncovered continuous molecular gradients and invasion trajectories indiscernible from individual 2D slices. The observed spatial interplay among CAFs, endothelial cells, and CAMs recapitulated key processes in tumor progression, including stromal remodeling, angiogenesis, and immune modulation, demonstrating that H2O can capture the multicellular dynamics of tumor invasion in three-dimensional space.

From a translational perspective, such 3D constructions provide new opportunities for histopathology-guided risk stratification and therapeutic targeting. By mapping invasive fronts, stromal niches, and immunomodulatory hotspots into three-dimensional space at low cost, H2O has potential to complement routine histopathology with molecular insights, ultimately informing prognosis and guiding precision oncology.

### H2O produces complementary spatial transcriptomics and spatial proteomics

While spatial transcriptomics offers a comprehensive window into cellular states, spatial proteomics provides a more immediate reflection of tissue functionality. Translating from histopathology to SP serves as an important complement to ST prediction and a key capability for general molecular prediction model. Extending H2O from ST to SP provides critical advantages. SP delivers a high-fidelity readout of cellular states that bypasses post-transcriptional buffering, reflecting disease pathology more accurately than ST. In addition to that, the targeted nature of SP facilitates the precise characterization of clinically actionable drivers, such as immune checkpoints, that directly govern therapeutic response.

To explore this possibility, we applied H2O to the Human Tumor Atlas Pilot Project (HTAPP) [65] from the Human Tumor Atlas Network (HTAN) project [66] which provides 8 paired samples of breast cancer profiled with MERFISH (ST) and CODEX (SP) (Fig. 6a). We first tested whether transcriptome-guided image embeddings could extend beyond RNA expression to inform protein-level distribution. With the eight samples containing histopathology-transcriptome-proteome data, we conducted a leave-one-out evaluation, training on seven samples with ST profiles and SP profiles, respectively, and testing on the held-out samples. To jointly model both data modalities, we introduced a parallel SP decoder, sharing an identical architecture with the original ST decoder. This dual-decoder framework enabled the model to simultaneously predict both ST and SP distributions from the same histopathological image input. Strikingly, we found that H2O predicted proteomics features more accurately than transcriptomics features in this experiment (Fig. 6b). We attribute this finding to the less sparse distribution of protein expression from the CODEX data. We generated spatial maps for key lineage-specific signatures, such as T-cell (*CD2, CD4, CD5*), B-cell (*CXCR5, CD38, CD40*), and macrophage (*HLA-DRA, HLA-B, MMP12*). These visualizations demonstrated a broad correlation between ST and SP, while also capturing platform-specific nuances in signature intensity and distribution (Fig. 6c,d).

**Fig. 6.**
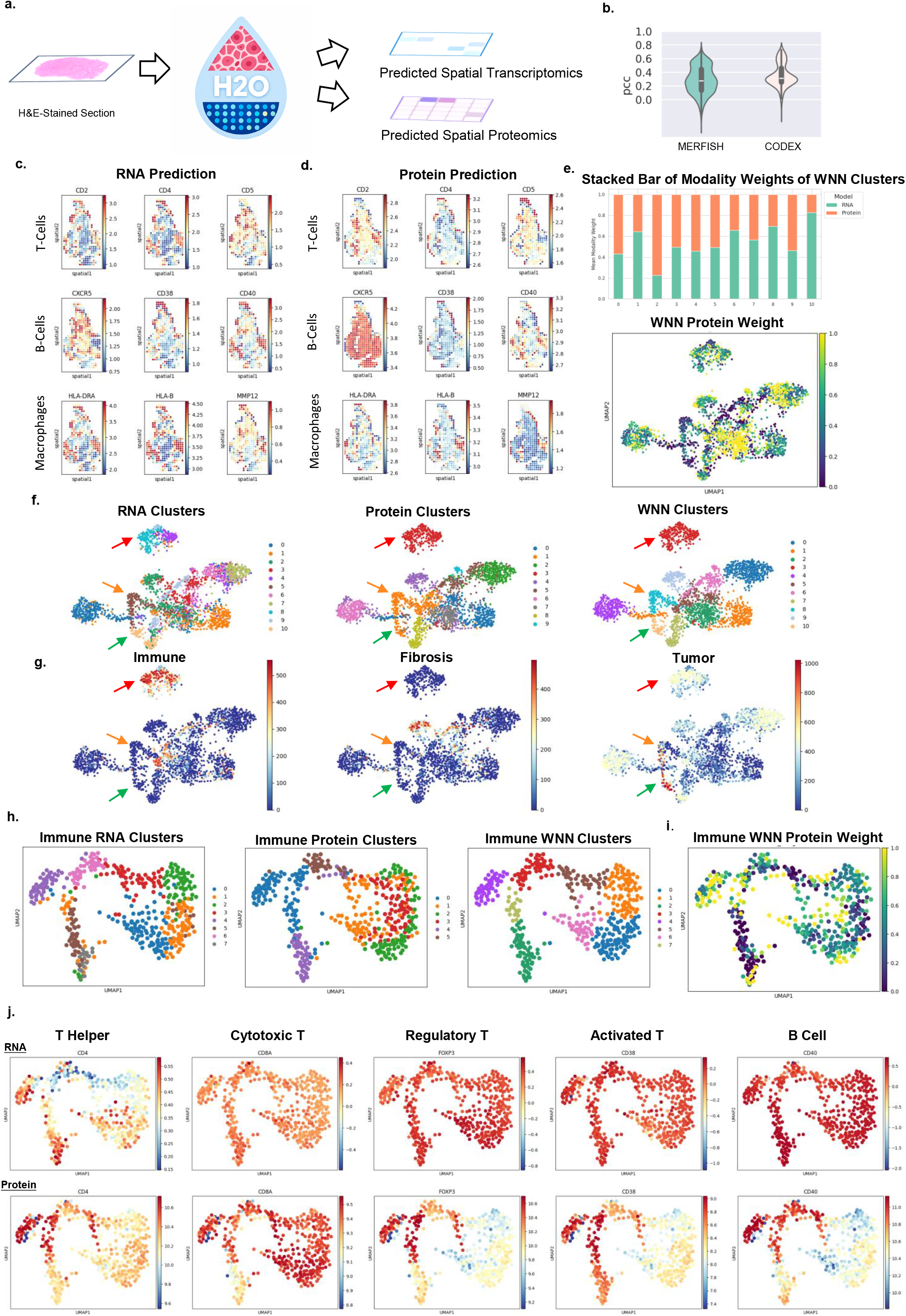
H2O produces complementary spatial transcriptomics and spatial proteomics. **a**, Overview of the workflow for applying H2O to the MERFISH and CODEX data from the HTAPP dataset. H&E images from consecutive sections are used to construct both ST and SP profiles using H2O. **b**, Average PCC metrics of eight paired MERFISH-CODEX samples evaluating H2O-predicted ST and SP with experiment-derived profiles, respectively. Performance of models trained on RNA (MERFISH) and protein (CODEX) were assessed and displayed separately, showing that transcriptome-guided training provides possibility for direct protein prediction from H&E images. **c-d**, H2O-predicted ST and SP maps of representative markers for T cells, B cells, and macrophages across selected sections. The predicted profiles reproduced both the spatial positioning of immune compartments and the expected tissue organization, demonstrating that H2O captured not only molecular signals but also their microanatomical context. **e**, Upper: Stacked bar plot displaying the relative RNA and protein modality weights across clusters derived from weighted nearest neighbor (WNN) integration followed by Leiden clustering. Lower: UMAP visualization of WNN embeddings of the eight samples colored by the protein modality weight. **f**, UMAP visualizations colored by Leiden clusters of RNA (MERFISH), protein (CODEX), and WNN (integration of both modality), respectively. **g**, UMAP plots illustrating three representative biological processes and cell-type signatures identified by the original HTAPP dataset in the WNN embedding space, including immune cells, fibrosis, and tumor-associated regions. **h**, Comparison of Leiden clusters derived from RNA (MERFISH), protein (CODEX), and integrated multi-modality (WNN) embeddings of immune cells. **i**, The weight of the protein modality in WNN clustering of immune cells, showing the contribution of protein signals in the integrated multi-modal clustering. **j**, Heatmap of immune-cell marker genes profiled with ST (upper row) and SP (lower row) predicted by H2O, showing the complementarity between the two modalities.

To investigate how the predicted RNA and protein modalities jointly contribute to downstream applications such as cellular clustering, we applied Seurat’s bi-modality integration framework (WNN) [67] followed by Leiden clustering. The resulting modality weights during clustering demonstrated contributions from RNA and protein features across clusters, revealing the complementarity and balance between the two modalities (Fig. 6e). Notably, in specific samples, proteomic signals contributed decisive information, as demonstrated by cluster-2 (green in the WNN clusters, Fig. 6f) resolved dominantly by proteomic features and not evident in transcriptomic space (Fig.6e,f). Fig. 6g highlights three cell-type signatures of representative biological processes identified by the original HTAPP dataset in the WNN embedding space, including immune cells, fibrosis, and tumor-associated regions, all of which are closely related to specific WNN clusters, demonstrating the reliability and practicality of the clustering results. Our results further demonstrated that the integration of predicted ST and SP signals refines tissue characterization beyond the capacity of single-modality profiling. Specifically, in the area indicated by the red arrow in Fig. 6f,g, WNN analysis identified tumor clusters that remained over-fragmented only by RNA profiling. While the orange and green arrows in Fig. 6f,g pointed out a distinct tumor sub-region and the cell type abundance label that were clearly distinguished in the WNN clusters but were obscured or poorly defined in the protein-only analysis. These observations underscored that harmonizing the two disparate signals enables a more robust and nuanced characterization of the complex regulatory landscapes within the tumor microenvironment.

Then, we particularly took the immune cells as an example for in-depth analysis. Comparison of Leiden clustering across RNA, Protein, and WNN embeddings of immune cells revealed that multimodal integration shows different results in cluster separability and biological interpretability (Fig. 6h). Visualization of the protein modality weight demonstrated the contribution of proteomic signals in WNN clustering (Fig. 6i). To further examine the WNN clusters of immune cells, we visualized the H2O-predicted expression of predefined markers in both ST and SP modalities within the joint clustering embedding space (Fig. 6j). The results highlighted the essential role of protein signals, which exhibited stronger heterogeneity and clearer patterns than the RNA modality, especially in regulatory T cells, activated T cells and B cells.

Together, our results demonstrate that H2O functions as a unified image-to-omics framework, generating spatially resolved transcriptomics and proteomics predictions directly from histology with high biological coherence. The integration of these two predicted modalities facilitates a synergistic characterization of tissue architecture, where the transcriptomics and proteomics landscapes provide complementary insights that transcend the limitations of single-modality profiling. By bridging histopathology with proteomic readouts, H2O provides a robust platform for inferring functional cellular states that are often inaccessible through transcriptomics alone, effectively translating morphological features into actionable proteomic insights.

## Discussion

In this study, we present H2O, a foundational framework that transcends simple morphological correlations by anchoring histopathological features into a high-dimensional molecular space via spatial omics guidance. Beyond outperforming existing benchmarks in predictive accuracy and cross-dataset generalization, H2O delivers superior biological resolution by recovering coherent molecular hierarchies and organized niches often invisible to standard histology. Specifically, through high-fidelity clustering and intercellular communication analysis, H2O successfully elucidated active signaling circuits, such as the immune modulators *MIF-CD44* and *MIF-CD74*, demonstrating its unique capacity to reconstruct functionally connected networks rather than isolated molecular markers.

The robustness of H2O was further underscored by its exceptional generality across diverse temporal, spatial, and multi-omics contexts. In developmental biology, H2O successfully preserved the intricate molecular trajectories of embryonic maturation, accurately capturing time-resolved gene expression programs from histological snapshots. Extending to three-dimensional landscapes, H2O generated volumetric molecular maps from serial sections of metastatic lymph nodes, delineating continuous invasion fronts and immune niches that remain invisible in 2D analysis. Beyond transcriptomics, the contrastive alignment paradigm of H2O successfully generalizes to spatial proteomics (SP), enabling high-fidelity RNA-protein profiling even from modality-limited samples. In breast cancer, this multimodal integration resolved functionally distinct cell states that were otherwise obscured in single-modality data. Collectively, these results demonstrate that H2O transcends specific biological contexts, providing a unified framework to decode complex tissue organization across dimensions and molecular layers.

A central challenge in spatial foundation models is reconciling the disparate biological scales across ST platforms, where identical pixel dimensions can represent anything from subcellular resolution to multi-cellular aggregates. To navigate this technical heterogeneity, H2O explicitly incorporates multi-scale supervision through two key mechanisms. First, by conditioning on platform-specific information such as spot diameter, H2O adaptively modulates its feature interpretation to match the specific resolution of different datasets. Second, by integrating neighboring patch context, H2O moves beyond local morphological cues to capture broader tissue architecture, such as vascular niches and immune gradients that govern spatially diffuse gene expression. Our findings underscore that the synergy between diameter-conditioned modulation and spatial context is essential for mitigating platform-specific variability, ensuring that image-based predictions remain biologically faithful across the diverse landscape of spatial omics technologies.

Despite these advances, several limitations remain. Although H2O leverages large-scale ST datasets for pretraining, coverage of rare tissues and underrepresented conditions is limited, which may constrain performance in specific contexts. Further expansion with single-cell atlases and bulk transcriptomics resources could improve zero-shot generalization. Moreover, H2O currently functions as a predictive framework rather than a generative one. Future integration with generative models may enable synthesis of full transcriptomics or proteomics maps at whole-slide scale.

Another important limitation is that our current implementation does not explicitly operate at cellular resolution. Instead, H2O predictions are derived from a fixed 224×224-pixel patch together with eight immediate neighbors, providing contextual but not truly single-cell-level information. While this patch-plus-context strategy effectively captures local and regional structure, it may miss molecular heterogeneity at the level of individual cells. Future work should explore multi-resolution learning with single-cell resolution as a basis, aggregating features across spatial scales, and directly integrating expression profiles from single-cell spatial multi-omics datasets once sufficient curated resources become available for foundation model training. Such improvements could enhance both the granularity and accuracy of predictions, ultimately bridging cellular and tissue-level organization within a unified framework.

In addition, while our extension of H2O to SP highlights the feasibility of translating histology into protein-level predictions, it remains limited. Unlike ST prediction, which benefits from pretrained encoders trained on large-scale ST datasets, SP translation in H2O relies only on a lightweight multilayer perceptron (MLP) without access to comparably rich pretraining. This restricts its ability to fully capture the complexity of protein regulation and pathway-level organization. Developing dedicated proteome-aware encoders or multimodal pretraining strategies that jointly leverage transcriptomics and proteomics datasets will be crucial to advance beyond proof-of-principle and achieve more reliable protein-level prediction from histology. Furthermore, expanding transcriptome-guided training to integrate additional omics layers, such as metabolomics, could establish multi-omics image foundation models with broader applications.

In summary, H2O establishes the concept of a spatial omics-guided image foundation model, in which histopathology image embeddings are trained under direct molecular supervision to produce spatial multi-omics prediction. This paradigm enables scalable, generalizable, and biologically interpretable representations of tissue organization, moving beyond morphology alone to recover conserved programs, dynamic transitions, and functional signaling networks. By transforming routine H&E slides into molecularly annotated maps, H2O pave the way for next-generation computational histopathology, bridging basic biology and clinical precision medicine.

## Methods

### Datasets

#### HEST-1k

We first fine-tuned scGPT on a subset of the HEST-1k dataset to generate scGPT-FT. Using the same subset, we trained H2O to obtain its pretrained checkpoint. For benchmarking against other methods, we trained H2O on predefined HEST-1k subsets, yielding task-specific models H2O-ccRCC and H2O-PRAD. Model generalizability was evaluated using the HEST-1k-LymphNodeIDC cohort as an independent test set, comprising samples that were strictly withheld from both the scGPT fine-tuning and H2O training stages.

The HEST-1k dataset contains over 60,000 recognized gene symbols, with each sample capturing only a subset of these genes due to differences in sequencing coverage, gene mapping and tissue diversity. To standardize feature space across samples, we identified the most variable genes within the dataset and selected the top 5,000. We then added genes sharing the same gene symbols with proteomic markers, resulting in a unified panel of 5,033 genes. This gene set was consistently used in both training and evaluation across all datasets.

#### OpenST-Metastatic Lymph Node

This dataset contains 19 serial sections (S2-S38) from a single metastatic lymph node, each profiled by ST and matched H&E images. We fine-tuned H2O-HEST-1k on section S2 and applied the model to predict expression across the remaining sections, enabling 3D construction of ST for the entire tissue.

#### Human Thymus Cell Atlas (HTSA)

HTSA provides Visium-based ST together with H&E images across fetal and early paediatric thymus samples spanning from 13 weeks post-fertilization to 2 years of age. At each time stage there are 2 or 4 samples except the 19-month-old stage and 19th week post-fertilization stage which each has only one sample. For both developmental stages, we used half of each time stage and the 19^th^ week post-fertilization stage sample as training. The other half samples and the 19-month-old stage sample were used as test samples.

#### HTAPP

The HTAPP dataset includes matched MERFISH transcriptomics, CODEX proteomics, and H&E images from the same samples. This multi-omics resource was used to assess the generality of H2O across ST and SP. We applied Leave-One-Out-Validation (LOOV) on all eight multimodal H&E-CODEX-MERFISH samples of the dataset.

### Model Training

We first fine-tuned scGPT on a subset of HEST-1k to adapt its transcriptomics representations to ST context. Then, we trained a Contrastive Language-Image Pretraining (CLIP) -based foundation model on the HEST-1k data pairs to align an in-house morphological FM with the fine-tuned scGPT (scGPT-FT), transforming the image FM into a bi-modal representation model that encodes both morphological and molecular information.

For each cell *i*, we combine (1) an image embedding from a DINOv2-trained ViT, (2) a neighborhood context aggregated by a 1D convolution across *K* adjacent patches, and (3) a conditioning signal from the spot diameter via FiLM to generate a representation. The resulting representation is used for contrastive alignment with a text embedding *h*^*i*^ and constructing the cell’s gene-expression profiles of the cell.

### Gene embeddings

#### Data preprocessing

ST expression data were represented as a cell-by-gene (or spot-by-gene) matrix, in which each element corresponds to the observed RNA abundance of a gene in a cell or spot. More generally, in single-cell RNA-seq (scRNA-seq), the element represents the read count of an RNA molecule, while it denotes chromatin accessibility of a peak region in scATAC-seq. Specifically, for ST data, each entry encodes the expression level of gene *j* ∈ {0, 1, …, G} in cell/spot *i* ∈ {0, 1, …, N}. We refer to this matrix as the raw count matrix.

We use the scGPT architecture with pretrained weights, followed by additional fine-tuning, as the gene encoder. The input to fine-tune scGPT consists of three components: (1) gene (or peak) tokens, (2) expression values, and (3) condition tokens. For each modeling task, gene tokens and expression values were derived from the raw count matrix X.

#### Gene tokens

We follow up scGPT’s scale and tokenize the ST expression data. Each cell/spot *i* is represented by a gene vector 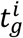 in *N*^*M*^:

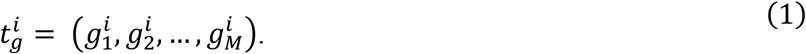

#### Expression values

The gene expression matrix X requires additional preprocessing before being used as input for the model. We applied log1p transformation through Scanpyto the raw matrices. For each gene *g* in cell/spot *i*, the expression value 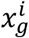 is defined as:

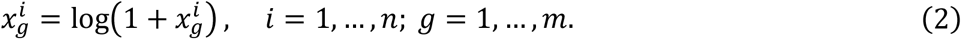

Therefore, the input matrix *X*^i^ can be defined as:

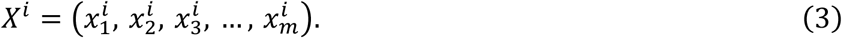

#### Condition tokens

The condition tokens encompass diverse meta-information associated with individual genes, such as perturbation tokens indicating perturbation experiment alterations. To represent position-wise condition tokens, we use an additional input vector that shares the same dimension as the input genes, denoted as:

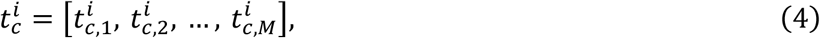

#### Embedding layers

We use the conventional embedding layers implemented in PyTorch for the gene tokens and condition tokens, denoted as emb_*g*_ and emb_*c*_, respectively, to facilitate the mapping of each token to a fixed-length embedding vector of dimension *D*. We use fully connected layers, denoted as emb_*x*_, for the binned expression values to enhance expressivity. This choice enables the modeling of the ordinal relation of gene expression values. Consequently, the final embedding *h*^*i*^ for cell *i* is defined as:

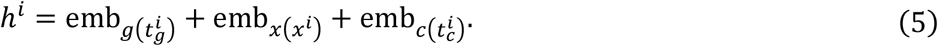

### Image embeddings

#### Image encoder and projection

The raw H&E image of cell *i* is encoded by the ViT into a 384-dimension vector and then projected to 512-dimension:

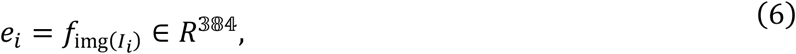

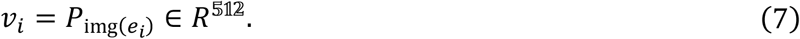

#### Neighborhood aggregation

In line with human visual processing of H&E images, which relies on contextual information from surrounding regions to infer details of a central location, we integrated neighborhood context during the training of each patch. For K nearest neighbor patches {*I*_i,1_, *I*_i,2_, …, *I*_*i,K*_}, we encode them with the same ViT and then stack the resulting embeddings along the neighbor axis, followed by applying 1D convolution (in_channels=384, out_channels=512, kernel_size=9) and Global Average Pooling (GAP) to obtain a 512-dimension. This procedure is defined as following formulas:

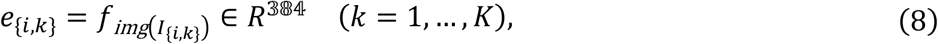

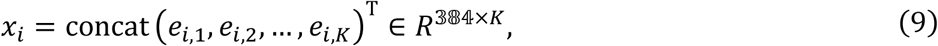

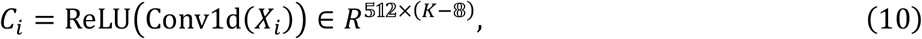

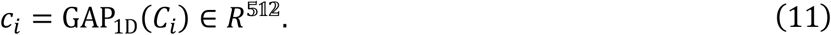

Both streams are layer-normalized:

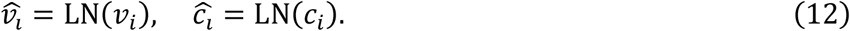

#### Diameter-conditioned FiLM

We employed Feature-wise Linear Modulation (FiLM), a conditioning mechanism that applies learned, input-dependent scaling and shifting to feature maps. By generating FiLM parameters from diameter embeddings, the model dynamically adjusted its representations according to spot size, thereby capturing scale-sensitive morphology-expression relationships and improving generalizability across platforms. All histopathology patches have a size of 224×224 pixels for foundation model training. Since the original whole slide images of the patches vary in resolution, the patches represent different scales of the true tissue size, thereby discarding absolute scale. However, because ST counts reflect the accumulation of transcripts over a defined physical area, the loss of scale may distort the link between morphology and total gene expression. In addition, spot diameters vary across platforms (e.g., Visium vs. Visium HD vs. imaging assays), introducing heterogeneity in both resolution and captured area. We applied FiLM module, which a diameter index *d*_*i*_ ∈ {1, … ,5} representing 2μm, 55μm, 100μm, 150μm or other diameter is embedded to 128-dimension and linearly mapped to FiLM parameters (γ_*i*_, β_*i*_) ∈ 512 × 512 shared for both streams. The formulas are defined as:

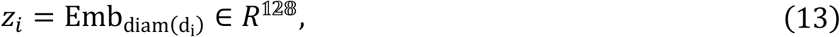

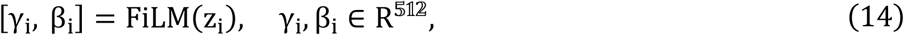

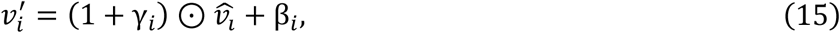

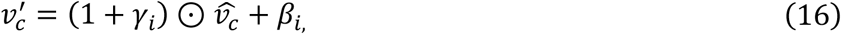

where 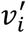 represents the image feature and 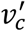 represents the contexture feature.

#### Concatenation and decoder

Finally, the two 512-dimension vectors are concatenated as *u*_*i*_ and decoded to the *H*-gene space (H = number of HVGs) through a decoder with 3-layer MLP, which is then denoted as *ĝ*_*i*_:

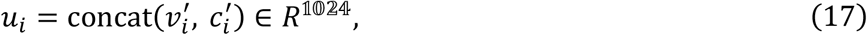

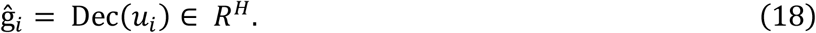

### Training objective

#### Fine-tuning Gene Encoder

We adopted scGPT as the backbone of our gene encoder. Unlike its original pretraining using discretized single-cell RNA-seq data, our task required the encoder to recover continuous gene expression values. To this end, we fine-tuned scGPT on real-valued expression matrices while retaining the original model architecture. The fine-tuning objective was to jointly reconstruct both gene identity and its associated continuous expression level, enabling the encoder to better capture quantitative variation in ST data.

#### Contrastive Loss

To align the two modalities, we adopt a contrastive objective based on a temperature-scaled dot product between image embeddings and text embeddings. In each mini-batch, paired image-text embeddings are regarded as positive examples, while all remaining cross-pairs within the batch serve as negative examples. To provide smoother targets, cosine similarity within each modality is additionally incorporated. The resulting contrastive loss is defined as:

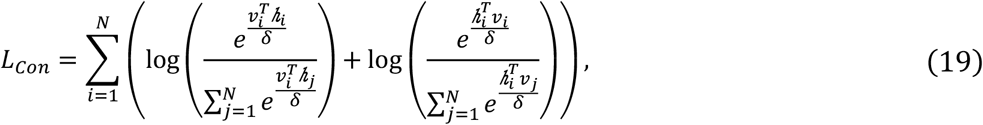

Where *v*_*i*_, *v*_*j*_ are patch image embeddings and *h*_*i*_, *h*_*j*_ are the ST embeddings for each cell/spot *i, j* in the batch.

#### Reconstruction Loss

The reconstruction objective was formulated as the mean squared error (MSE) between the predicted expression profiles *ĝ*_*i*_ and the original experimental profiles *g*_*i*_:

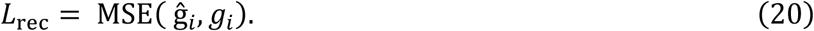

#### Joint Loss

The overall training objective combines the reconstruction loss with the contrastive alignment loss:

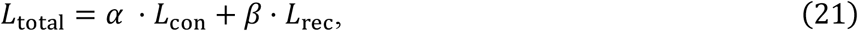

where *α* and *β* are fixed weighting coefficients that balance the two terms.

### Evaluation Metrics

To assess the performance of H2O, we employed four complementary evaluation metrics: Pearson correlation coefficient (PCC), Spearman’s rank correlation coefficient (SRCC), concordance correlation coefficient (CCC), and root mean squared error (RMSE).

#### Pearson Correlation Coefficient (PCC)

For each gene i in the predicted gene set, we computed the PCC metric, denoted as PC*C*_*i*_ between the observed expression vector *G*_*i*_ and the predicted expression vector 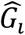:

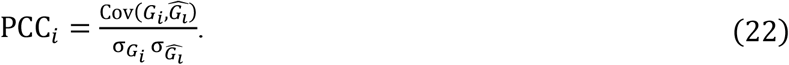

#### Spearman’s Rank Correlation Coefficient (SRCC)

To assess monotonic relationships between the observed and predicted expression independent of absolute values, we calculated SRCC based on the rank-transformed vectors R(*G*_*i*_) and 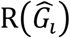:

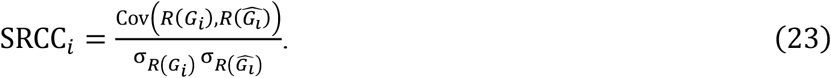

#### Concordance Correlation Coefficient (CCC)

To quantify numerical accuracy of the predictions, we computed CCC as:

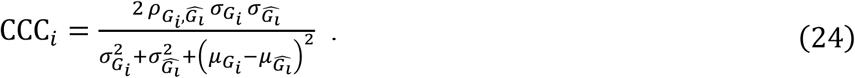

#### Root Mean Squared Error (RMSE)

Finally, the prediction error was quantified by RMSE:

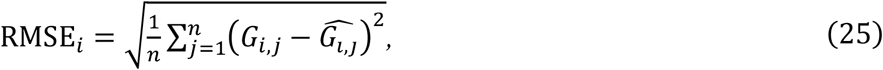

where *i* is the number of cells/spots.

### Benchmark

To evaluate the expression generation ability of H2O from histopathology images, we benchmarked it against four state-of-the-art methods, including OmiCLIP, HisToGene, BLEEP, and DeepPT. Together with H2O, these five approaches were systematically trained and assessed on the two HEST-1k subsets, including HEST-1k-ccRCC and HEST-1k-PRAD, using the four evaluation metrics defined above. To further examine the generalizability of these methods, we additionally tested them on the additional HEST-1k-IDC-LymphNode dataset, using models pretrained on the HEST-1k-ccRCC and HEST-1k-PRAD benchmarks.

#### HisToGene

HisToGene was originally designed to operate on 112×112 image patches. To ensure fair comparison with other methods, where the patch size directly influences the amount of information conveyed, we adapted HisToGene to use 224×224 patches while keeping the rest of the architecture unchanged.

#### BLEEP

BLEEP leverages contrastive learning to construct a low-dimensional joint embedding space from a reference dataset containing paired image and gene expression profiles at micrometer resolution. It employs a ResNet50 pretrained on ImageNet as the image encoder and a fully connected network as the expression encoder. We adopted the preprocessing pipeline and architectural framework of BLEEP. The training set for training or fine-tuning other methods are used as the reference dataset for BLEEP.

#### DeepPT

DeepPT represents the first stage of the ENLIGHT-DeepPT framework, which predicts genome-wide tumor mRNA expression directly from histopathology slides. In this study, we adapted DeepPT for application to ST data. DeepPT employs a ResNet-50 backbone pretrained on ImageNet as the image encoder. To reduce the dynamic range of expression values and mitigate batch- and experiment-specific discrepancies in library size, we followed the normalization scheme described in the original DeepPT paper.

#### OmiCLIP

OmiCLIP is a visual–omics foundation model. In our framework, OmiCLIP Align and OmiCLIP Retrieve is used for H&E image-to-transcriptomics retrieval, while Loki PredEx is applied to predict gene expression from image patches using the reference ST-image data for retrieve. On the HEST1K-ccRCC and HEST1K-PRAD benchmarks, we predict expression for the held-out samples using the remaining samples within each benchmark as reference data.

## Data availability

All datasets used in this study are publicly available and were downloaded from the following links. The raw data can be download from the following link: HEST-1k for H2O training and testing: https://huggingface.co/datasets/MahmoodLab/hest. HTSA dataset for human development analysis: https://cellxgene.cziscience.com/collections/fc19ae6c-d7c1-4dce-b703-62c5d52061b4. OpenST for 3D transcriptome reconstruction: https://rajewsky-lab.github.io/openst/latest/examples/datasets/. HTAPP for multi-omics: The preprocessed (previously spatially aligned multi-omics expression and images) data can be downloaded at: https://singlecell.broadinstitute.org/single_cell/study/SCP2702/htapp-mbc; Other raw material of HTAPP can be discovered in the HTAN portal: https://humantumoratlas.org/.

## Code availability

The open-source implementation of H2O, detailed tutorials and a demo are available in GitHub at https://github.com/TencentAILabHealthcare/H2O.

## Acknowledgements

This work is approved by the funding from China State Key Basic Research Program Grants (2024YFA1307200), National Natural Science Foundation of China (62402473, 62271465), and Suzhou Basic Research Program (SYG202338).

